# PertAdapt: Unlocking Single-Cell Foundation Models for Genetic Perturbation Prediction via Condition-Sensitive Adaptation

**DOI:** 10.1101/2025.11.21.689655

**Authors:** Ding Bai, Le Song, Eric P. Xing

## Abstract

Single-cell foundation models (FMs) pretrained on massive unlabeled scRNA-seq data show strong potential in predicting transcriptional responses to unseen genetic perturbations. However, existing approaches insufficiently transfer pretrained knowledge and overlook the imbalance between perturbation-sensitive and insensitive genes, yielding only marginal improvements over non-pretrained baselines. To address these limitations, we introduce PertAdapt, a framework that unlocks FMs to accurately predict genetic perturbation effects via integrating a plug-in perturbation adapter and an adaptive loss. The adapter employs a gene-similarity-masked attention mechanism to jointly encode perturbation conditions and contextualized representations of unperturbed cells, enabling more effective knowledge transfer. To better capture differential expression patterns, the adaptive loss dynamically reweights perturbation-sensitive genes relative to global transcriptomic signals. Extensive experiments across seven perturbation datasets, including both single- and double-gene settings, demonstrate that PertAdapt consistently outperforms non-pretrained and FM baselines. Moreover, PertAdapt demonstrates strong capacity for modeling multiplexed gene interactions, generalizing in limited-data regimes, and maintaining robustness across backbone sizes.

**Availability:** Code and data are available at https://github.com/BaiDing1234/PertAdapt.

## 1 Introduction

Many single-cell RNA sequencing (scRNA-seq) datasets of genetic perturbations have been generated through CRISPR-based perturbational screens, enabling systematic exploration of cellular responses to diverse perturbations [1, 2, 3, 4, 5, 6, 7]. Understanding these transcriptional responses is critical to deciphering the principles of gene regulation, while experimental approaches such as Perturb-seq and large-scale CRISPR screens remain constrained by cost, scalability, and feasibility, particularly when extending to unseen perturbations [1, 2, 8]. These limitations highlight the need for computational models capable of generalizing beyond measured conditions to predict cellular responses to novel perturbations [9, 10, 11, 12, 13, 14, 15], offering a scalable route toward mapping regulatory programs at a resolution unattainable by experiments alone.

Recently, Transformer-based single-cell foundation models (FMs) have emerged, trained on large-scale unlabeled single-cell transcriptomic datasets to solve various downstream tasks [16, 17, 18, 19, 20, 21, 22, 23]. Several of these models, such as scFoundation, scGPT, GeneCompass, and AIDO.Cell, explicitly include predicting perturbational effects as a key downstream task [18, 19, 20, 21, 23]. Others, including scBERT, Geneformer, and UCE, are not initially designed for perturbation prediction but can potentially be repurposed for this task [16, 17, 22, 24]. Despite their success in tasks such as cell-type clustering and classification, these foundation models have shown only limited improvement over the previous state-ofthe-art (SOTA) method GEARS [14] in the task of genetic perturbation effects prediction, particularly in their own reported benchmarks. Furthermore, independent evaluations indicate that, when tested under alternative benchmarks or data splits, they do not yet consistently outperform even simple linear regression approaches [24].

There are two major limitations in how current single-cell FMs are applied to predict transcriptional responses to genetic perturbations. **First**, the knowledge acquired from massive unlabeled datasets is not effectively transferred to downstream perturbation tasks. In natural language processing, pretrained language models achieve SOTA results largely through transfer learning strategies such as adapters [25], prefix tuning [26], and LoRA [27], which preserve pretrained representations while enabling efficient task-specific adaptation [28, 29, 30]. In contrast, most single-cell FMs directly attach either simple linear layers [20] or existing perturbation model GEARS [14, 18, 19, 21, 23], without adequately addressing the unique challenges of perturbation prediction, including large model scale, task-specific distribution shifts, and biological heterogeneity. **Second**, current approaches overlook the extreme imbalance between perturbation-sensitive and insensitive genes. Existing methods typically treat all readout genes equally in the loss computation, using either the direct mean squared error over all genes [20] or the GEARS loss [14, 18, 19, 21, 23]. Yet, for a given perturbation, only a very small subset (often fewer than 20) of the ∼ 20,000 readout genes exhibits distinct responses. This imbalance dilutes gradient updates across mostly uninformative targets, limiting effective optimization of the model. Together, these issues restrain the ability of current FMs to fully exploit pretrained knowledge for accurate perturbation effects prediction.

To overcome these limitations, we propose **PertAdapt**, an FM-based method for predicting transcriptional responses to genetic perturbations that integrates a plug-in perturbation adapter with an adaptive loss. To address the first limitation, we introduce a perturbation-conditional adapter that enables effective transfer of knowledge from pretrained FMs on unlabeled data to labeled perturbation conditions. Inspired by NLP adapter architectures, our perturbation adapter introduces a biologically specialized design that incorporates gene-level functional structure. Specifically, through an attention mask derived from gene-wise functional similarities, the adapter employs a masked multi-head attention [31] to jointly encode perturbation conditions and FM representations of unperturbed cells, obtaining comprehensive post-perturbation cell states. We apply this plug-in adapter to two backbone FMs, scFoundation and AIDO.Cell [19, 23], both of which encode full-gene expression profiles. To address the second limitation, we design an adaptive loss that accounts for the imbalance between perturbation-sensitive and insensitive genes. Rather than uniformly weighting all readout genes, the adaptive loss dynamically adjusts the contribution of sensitive genes relative to the background, prioritizing the learning of meaningful differential responses while preserving consistency across the transcriptome.

PertAdapt demonstrates substantial improvements over prior approaches, including simple baselines [24], non-pretraining deep-learning methods [14], and native applications of pretrained foundation models [19, 20, 21, 23], across benchmarks and datasets. In contrast to previous studies that typically relied on only two or three datasets, our evaluation spans seven diverse collections, providing a more comprehensive assessment of model generalization. These datasets cover distinct biological settings and were selected to probe three key aspects of performance: (*i*) prediction on multiple unseen gene perturbations, tested using the *Norman* dataset with 125 double-gene perturbations [2]; (*ii*) scalability across large single-gene perturbation screens in different cell lines, including *RPE1, K562, HepG2*, and *Jurkat* [6, 7], each comprising more than 1,000 perturbations; and (*iii*) robustness under limited-sample conditions, assessed using the *Adamson* dataset with 78 perturbations and the *Dixit* dataset with 20 perturbations. These extensive experiments demonstrate that PertAdapt effectively leverages pre-trained foundation models to achieve more accurate predictions of post-perturbation gene expression profiles and multiplexed interaction behaviors than existing approaches. In addition, our analyses reveal a scaling trend when applying PertAdapt to foundation models of varying sizes, highlighting its robustness across backbones with different capacities.

## 2 Methods

### 2.1 Problem setting

PertAdapt predicts single-cell gene expressions for post-perturbation cells given specific perturbed gene(s). Formally, the set of perturbed genes, also called a gene-perturbation condition, is denoted as *c* ⊆ {1, …, *N*} representing a subset of *N* gene indices. Within the datasets, each perturbation *c* is associated with *n*_*c*_ cells represented as {**ŷ**_*c*_ ∈ ℝ^*N*^}, where *n*_*c*_ is the number of perturbed cells under the condition *c*. The objective is to predict *n*_*c*_ post-perturbation gene expressions {ŷ_*c*_ ∈ ℝ^*N*^}. A perturbation condition is termed *control* when it is empty, meaning the cells under this condition are unperturbed normal cells, and the unperturbed expressions are considered as known entities denoted by {**x**^*i*^, *i* = 1, …, *n*_control_}.

Following prior work [14, 19], we pre-process each dataset separately using random pairing, where each dataset contains cells from one particular cell line. For every perturbed cell 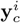 where *i* = 1, …, *n*_*c*_ for each condition *c*, a control cell 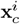 is uniformly randomly sampled from all unperturbed cells in pairs with the perturbed cell. In training and testing, models predict the perturbed cell 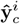 by inputting the paired control sample 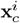 and the condition *c*. In this experimental setting, we trained and tested our model on each dataset separately, as shown in the Results Section.

### 2.2 Model architecture

We propose the perturbational adapter, a plug-in architecture for pretrained single-cell foundation models. As illustrated in Fig. 1, we employ a frozen pretrained single-cell foundation model to encode unperturbed expressions **x**_*c*_, and a perturbation condition encoder trained from scratch to encode the set of perturbed genes *c*, following prior work [14, 19, 23]. These two encodings are merged and fed into our perturbation adapter, which generates condition-wise and contextualized representations of perturbed expressions. The adapter integrates the encodings through masked multi-head attention, where the mask is derived from functional gene-gene similarities computed using Gene Ontology. Finally, the perturbed representations are passed through multi-layer perceptrons (MLPs) to produce post-perturbation predictions **ŷ**_*c*_. Formally, denoting the whole model as **g**_*θ*_, the output can be written as: **ŷ**_*c*_ = **g**_*θ*_(**x**_*c*_, *c*).

**Figure 1:**
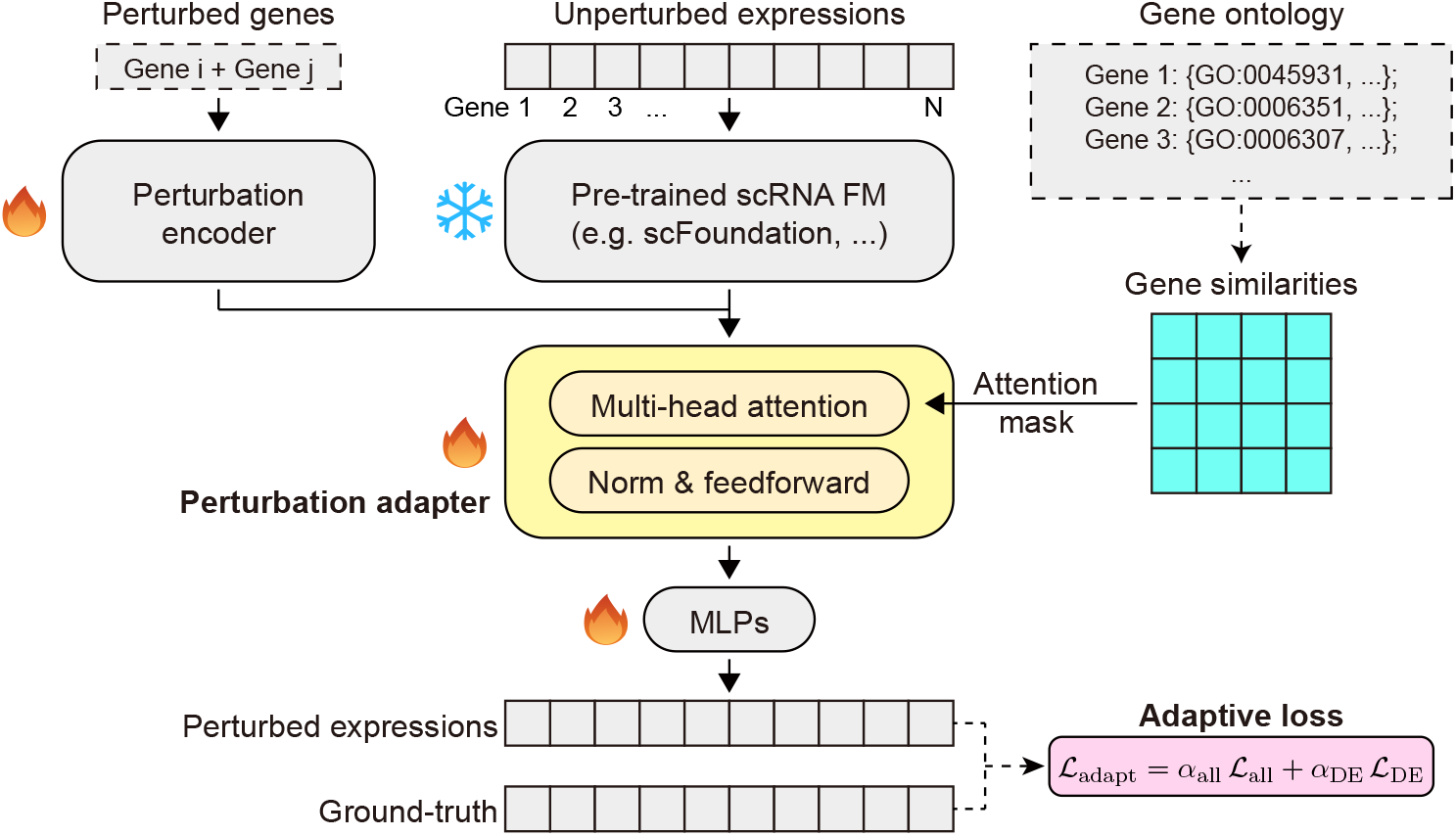
The architecture of the proposed PertAdapt. The model takes as input (i) the full-length gene expression vector of an unperturbed cell and (ii) its perturbation condition, represented as a set of perturbed genes. The unperturbed expression profiles are encoded by a pretrained single-cell foundation model (e.g., scFoundation, AIDO.Cell), while the perturbation condition is encoded by a perturbation encoder (here we use the GEARS encoder for single- or multi-gene perturbations). These two representations are integrated within the multi-head attention layer of our perturbation adapter, where a gene-gene similarity matrix derived from Gene Ontology (GO) terms is applied as the attention mask. The resulting embeddings are then passed through multi-layer perceptrons (MLPs) to generate predicted perturbed expression profiles. Model training is guided by our adaptive loss, dynamically weighting reconstruction losses on all genes ℒ_all_ and on perturbation-sensitive (i.e., differentially expressed, DE) genes ℒ_DE_, where the weights *α* are adjusted per batch. During training, only the added modules (the perturbation encoder, the adapter, and the MLPs) are updated, while the parameters of the pretrained foundation model remain frozen.

#### 2.2.1 Backbone FM and perturbation encoder

In our framework, the pretrained foundation model (FM) takes a control cell **x**_*c*_ ∈ ℝ^*N*^ as input and produces full-length contextualized representations *F* (**x**_*c*_) ∈ ℝ^*N ×D*^, where *D* denotes the latent dimensionality. Existing single-cell foundation models differ in the coverage of readout genes. Models such as Geneformer [17], scGPT [20], UCE [32] and GeneCompass [21] typically output representations for a cell-specific subset of genes (fewer than 3,000). In contrast, models like scBERT [16], scFoundation [19], and AIDO.Cell [23] generate encodings for the full set of ∼ 20,000 readout genes, with the sequence length fixed across all cells. Since our perturbation adapter requires the whole gene-spectrum representations, we adopt scFoundation and AIDO.Cell as backbone models, using their publicly available pretrained weights kept frozen, and train and evaluate our adapter on each of them separately.

The perturbation encoder is adopted from GEARS [14], a graph neural network (GNN)-based model originally designed to predict post-perturbation expression changes. We reuse its GNN encoder to embed perturbation conditions into the latent space and denote this encoder as **p**. For any perturbation condition *c*, the corresponding embedding is **p**(*c*) ∈ ℝ^*D*^. In the case of a double-gene perturbation *c* = {*j*_1_, *j*_2_}, the encoding is obtained by summing the embeddings of the two single-gene perturbations:**p**({*j*_1_, *j*_2_}) = **p**({*j*_1_}) + **p**({*j*_2_}). The perturbation encoder is trained from scratch.

#### 2.2.2 Perturbation adapter

The perturbation adapter consists of a masked multi-head self-attention (MHA) layer followed by layer normalization (LN) and feedforward (FF) networks. The input query is given by the contextualized expression encodings with the perturbation embedding added to each row: *Q* = LN *F* (**x**_*c*_) + **1**_*N*_ **p**(*c*)^⊤^, where LN denotes the layer normalization of the input, and **1**_*N*_ denotes a length-*N* vector of ones, ensuring that the perturbation encoding **p**(*c*) is added row-wise. The key and value are identical to the query, i.e., *K* = *V* = *Q* ∈ ℝ^*N* × *D*^. Then the output of the masked multi-head attention is:

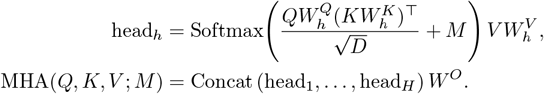

Here, matrices 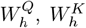 and 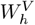 denote the projection weights of the *h*-th head for query, key, and value, respectively, and the matrix *W*^*O*^ is the output projection. All these *W*’s are trainable parameters. The attention mask *M* ∈ ℝ^*N ×N*^ is a pre-defined knowledge matrix encoding gene-wise functional similarities, as illustrated in Fig. 2a. Specifically,

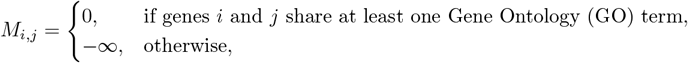

such that the attention weights between genes without any functional overlap are completely masked. GO is a curated hierarchical vocabulary that standardizes the representation of gene functions across biological processes, cellular components, and molecular functions [33, 34, 35]. Among various sources of prior gene-gene relationships, we selected GO because it provides the broadest coverage for our readout space, annotating 18,632 of the 19,264 genes. On average, each gene shares at least one GO term with 1,563 other genes (*≈* 8.1% of readout genes, detailed in Appendix Fig. A1), yielding a biologically meaningful yet sufficiently sparse topology for the attention mask. This prior structure is incorporated into the multi-head attention to bias information flow toward functionally related genes.

**Figure 2:**
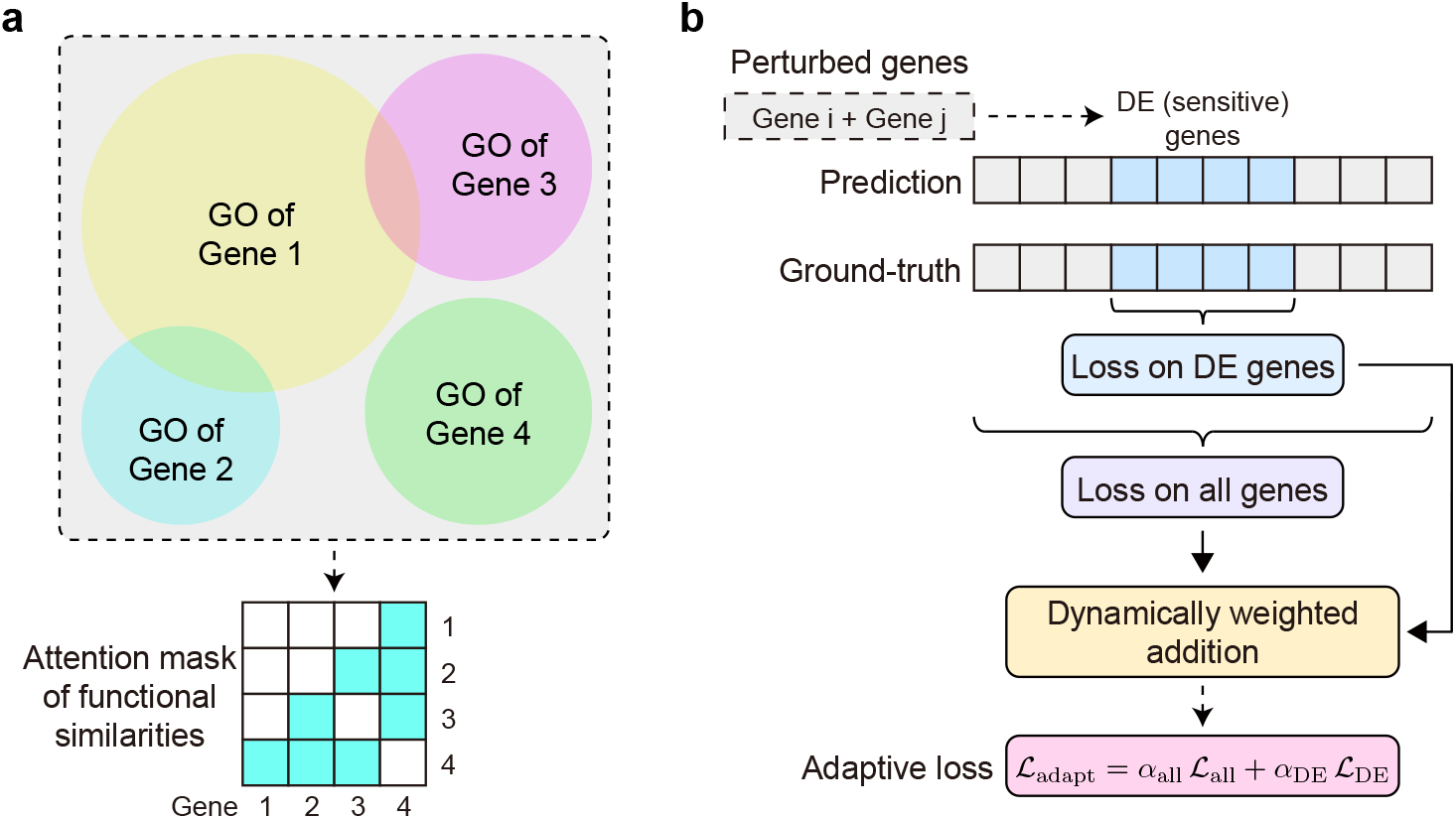
Method illustration of (a) the gene-similarity attention mask in the perturbation adapter and (b) the adaptive loss. **a**, The attention weight between two genes is masked (set to −∞) if the gene pair shares no common Gene Ontology (GO) term, otherwise remains unchanged. **b**, For a given perturbation condition, the set of perturbation-sensitive, i.e., the top 20 differentially expressed (DE) genes, is specified, and the adaptive loss is the dynamically weighted summation of the losses on all genes and on the DE genes.

The output of the masked multi-head attention, MHA(*Q, K, V*; *M*), is then input to the Norm & Feedforward structure of a Transformer encoder layer [31]. Specifically, it is first passed through a residual addition with the input of the MHA, followed by layer normalization. The normalized representations are then fed into a position-wise two-layer feedforward network with a ReLU activation, followed again by residual addition and layer normalization. The output contextualized representations, *E* ∈ ℝ^*N* × *D*^ that are further processed by subsequent MLP layers for final prediction.

#### 2.2.3 Final MLPs

The final layer consists of *N* independent MLP heads, each mapping the post-perturbation representation of a gene to a scalar value that is added to its unperturbed expression. Formally,

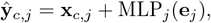

where MLP_*j*_ denotes the MLP associated with the *j*-th gene, which maps a *D*-dimensional input to a scalar, and **e**_*j*_ ∈ ℝ^*D*^ is the *j*-th row of the post-perturbation encodings *E* produced by the perturbation adapter. We then optimize the adaptive loss between **ŷ**_*c*_ and the ground-truth **y**_*c*_, updating all trainable components of the model, including the perturbation encoder, the perturbation adapter, and the output MLPs.

### 2.3 Adaptive loss

Apart from the model architecture of the perturbation adapter, we also introduce an adaptive loss designed to dynamically balance the reconstruction error between genes sensitive to perturbations and all readout genes. During training, a vector of perturbed expressions **ŷ**_*c*_ = **g**_*θ*_(**x**_*c*_, *c*) is predicted by the model and compared with the ground-truth perturbed cell **y**_*c*_. As illustrated in Fig. 2b, the loss combines the mean squared error (MSE) over all genes, ℒ_all_, with that over the selected top *k* differentially expressed (DE) genes, ℒ_DE_, in a dynamic, data-driven manner, as shown below.

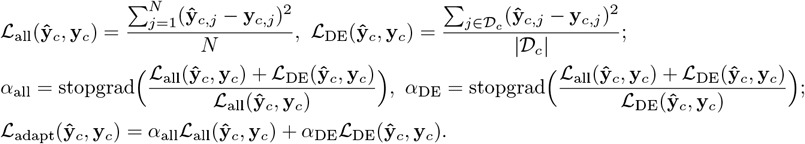

Here, the subscript *j* denotes the *j*-th readout gene within a single-cell expression vector, *N* is the number of all genes, and 𝒟_*c*_ is the set of top *k* DE genes under perturbation condition *c* (*k* is a tunable hyperparameter). We apply a stop-gradient operation (denoted as stopgrad(·) and implemented in practice with detach in PyTorch) to *α*_all_ and *α*_DE_, ensuring that the weighting coefficients are treated as constants during backpropagation. This prevents gradients from flowing through the adaptive weights, so that the coefficients reflect only the instantaneous relative magnitudes of the MSE losses. These adaptive weights are updated dynamically at each forward pass but introduce no additional trainable parameters. Note that the denominators never vanish in practice, as training loss does not converge to exactly zero. The batch loss is then computed as the sum of the above loss across all prediction/ground-truth pairs and optimized using Adam [36].

### 2.4 Experimental setup

#### 2.4.1 Evaluation metrics

The evaluation metrics for each perturbation condition are the mean squared error (MSE) and the Pearson correlation coefficient, computed between the predicted post-perturbation pseudobulk expressions (the mean expression of each gene across all cells under the same perturbation condition) and the corresponding ground-truth (detailed in Appendix Section B). In the next section, we present the MSE and Pearson correlation scores on the top 20 DE genes, while results for other numbers of top DE genes or all genes are reported in Appendix Section D.

#### 2.4.2 Baselines and backbones

We evaluated 7 baseline methods and compared their performance against PertAdapt. First, we included 4 pretrained single-cell foundation models (FMs) that originally reported perturbation-effect prediction results: scGPT [20], GeneCompass [21], scFoundation [19], and AIDO.Cell [23]. We inferred their performances using their native finetuning procedures and public pretrained weights. In addition, we included two task-specific approaches, GEARS [14] and a PCA-based linear model [24], both without pretraining. Finally, we included *No-Change* [24] which directly uses the input unperturbed expressions as the output. For all baseline models, we strictly follow their original architectural and optimizer hyperparameters as reported in their papers. To ensure each baseline is sufficiently trained, we extend their training epochs with a unified early-stopping strategy, allowing every method to fully converge under comparable conditions.

For PertAdapt backbone FMs, we used scFoundation and AIDO.Cell, which provide full-spectrum representations for the 19,264 genes. For scFoundation, we used its only released 100M-parameter checkpoint. For AIDO.Cell, we examined scaling using the 3M, 10M, and 100M checkpoints, and selected the best 100M model as representative for comparison. Following the perturbation downstream methods of scFoundation and AIDO.Cell, we adopt a graph convolutional network (GCN) built on a GO-derived graph, as in GEARS, and train its weights from scratch as the backbone perturbation encoder. Backbone hyperparameters are listed in Appendix Section C.

#### 2.4.3 Implementation and datasets

PertAdapt was implemented using the PyTorch library in Python. The hyperparameters of PertAdapt were configured as follows: the number of DE genes in the adaptive loss is *k* = 20, the embedding dimension was set to *D* = 512, the number of attention heads to *H* = 8, and the head dimension to 64 in the perturbation adapter (grid search detailed in Appendix Section C). We trained the model for up to 20 epochs with early stopping based on the validation loss, using Distributed Data Parallel (DDP) on 4 GPUs. Each GPU processed a batch of 4 samples, and gradients were accumulated for 2 steps, resulting in an effective batch size of 4*×* 4*×* 2 = 32. We used the Adam optimizer with a learning rate of 10^−3^.

Model training and evaluation for all baselines and PertAdapt was conducted on 7 datasets: Norman, Replogle RPE1, Replogle K562, Nadig HepG2, Nadig Jurkat, Adamson, and Dixit [2, 6, 7, 37, 1]. Among them, the Norman dataset contains ∼ 200 double-gene perturbations, the Replogle and Nadig datasets each contain over 1,000 single-gene perturbations, while the Adamson and Dixit datasets each include fewer than 80 single-gene perturbations. More dataset details are provided in Appendix Section A. In each dataset, cells related to 25% of the perturbations were in the test set while the cells of the remaining 75% perturbations, including unperturbed cells, were in the train-validation set. The train-validation set was further split to 90% for training and 10% for validation. For each dataset, we randomly generated five independent train/validation/test sets. The model was trained and evaluated on each split, and the average performance across the five runs was reported.

## 3 Results

In this section, the best results in each table are highlighted in **bold**, and the second-best results are <u>underlined</u>. The symbols “↑“ and “↓“ after each metric indicate that higher or lower values are better, respectively. All reported metrics (MSE and Pearson) are evaluated on the top 20 differentially expressed (DE) genes, while results for different numbers of DE genes or all genes are in Appendix Section D.

### 3.1 Modeling double-gene perturbation and gene interactions

PertAdapt is adept at predicting transcriptional responses to double-gene perturbations, with comparative results on the Norman dataset summarized in Table 1. To evaluate perturbation-level generalization, we split the data by perturbation condition *c*, ensuring that all cells under the same condition fall entirely into either the training/validation set or the test set. For double-gene perturbations, there are three separate scenarios based on whether each gene in the tested pair has been seen during training: seen 0/2 (neither gene seen), seen 1/2 (only one gene seen), and seen 2/2 (both genes seen, but the combined perturbation is still unseen during training). Across all evaluation scenarios, PertAdapt consistently outperforms prior baseline models, regardless of the chosen backbone. In the most challenging out-of-distribution (OOD) setting, where neither test gene is exposed during training (the seen 0/2 case), PertAdapt achieves the lowest and second-lowest mean squared error (MSE) with its two backbone variants, yielding improvements of approximately 11.5% and 11.2% over the corresponding native methods of scFoundation and AIDO.Cell, respectively. The largest gain over previous methods is observed in the seen 2/2 scenario, where PertAdapt with the scFoundation backbone reduces MSE by 42.6% and increases the Pearson correlation by 2.9%. Additional results in Appendix Section D further confirm the superiority of PertAdapt, demonstrating consistent improvements over most baselines when evaluating different numbers of DE genes or all genes on the Norman dataset.

**Table 1:** Evaluation of PertAdapt and baselines on the Norman dataset. We report the results on the overall scenario (all tested perturbations) and three training-exposure scenarios (seen 0/2, seen 1/2 and seen 2/2 perturbations).

| Model | Overall |  | Seen 0/2 |  | Seen 1/2 |  | Seen 2/2 |  |
| --- | --- | --- | --- | --- | --- | --- | --- | --- |
|  | MSE↓ | Pearson↑ | MSE↓ | Pearson↑ | MSE↓ | Pearson↑ | MSE↓ | Pearson↑ |
| No-Change | 0.380 | 0.726 | 0.443 | 0.538 | 0.413 | 0.679 | 0.377 | 0.769 |
| Linear Model | 0.231 | 0.780 | 0.336 | 0.561 | 0.237 | 0.745 | 0.122 | 0.877 |
| GEARS | 0.229 | 0.770 | 0.240 | 0.600 | 0.245 | 0.732 | 0.158 | 0.849 |
| scGPT | 0.221 | 0.765 | 0.231 | 0.599 | 0.228 | 0.739 | 0.141 | 0.844 |
| GeneCompass | 0.195 | 0.773 | 0.226 | 0.605 | 0.221 | 0.729 | 0.144 | 0.837 |
| scFoundation | 0.210 | 0.776 | 0.234 | 0.603 | 0.211 | 0.746 | 0.128 | 0.859 |
| AIDO.Cell | 0.192 | 0.781 | 0.232 | 0.611 | 0.201 | 0.753 | 0.124 | 0.870 |
| scFoundation+Ours | **0.156** | **0.789** | <u>0.207</u> | **0.614** | **0.156** | <u>0.754</u> | **0.070** | **0.902** |
| AIDO.Cell+Ours | <u>0.159</u> | <u>0.786</u> | **0.206** | <u>0.612</u> | <u>0.160</u> | **0.756** | <u>0.080</u> | <u>0.890</u> |

Furthermore, to characterize the gene interaction (GI) types of double-gene perturbations, we follow [2] and adopt two quantitative measures: the magnitude and the model fit scores, for GI types: synergy and neomorphism, respectively. A larger magnitude for a double perturbation {*i, j*} indicates stronger synergy between *i* and *j*. A lower model fit suggests that the effects of individual perturbations {*i*} and {*j*} are less relevant to the combined response of {*i, j*}, indicative of neomorphic behavior (detailed in Appendix Section B). We further evaluate all models by computing the MSE between the predicted and ground-truth magnitude and model fit scores. As shown in Fig. 3, PertAdapt achieves the lowest MSE for both metrics that indicate GI tendencies: magnitude for synergy and model fit for neomorphism, demonstrating superior accuracy in recovering the underlying signals of double-gene perturbational GI types. These improvements suggest that PertAdapt provides a more faithful reconstruction of the combined perturbational effects, thereby offering more reliable estimates of interaction tendencies indicative of synergy or neomorphism. Between the two top-performing variants, scFoundation+PertAdapt achieves lower magnitude-prediction MSE (Fig. 3) and better seen 2/2 performance (Table 1) than AIDO.Cell+PertAdapt, while the two show comparable results on the remaining metrics. Since both outperform all other baseline methods across the board, we conclude that scFoundation+PertAdapt achieves the strongest overall performance on the Norman dataset among all methods evaluated in this study.

**Figure 3:**
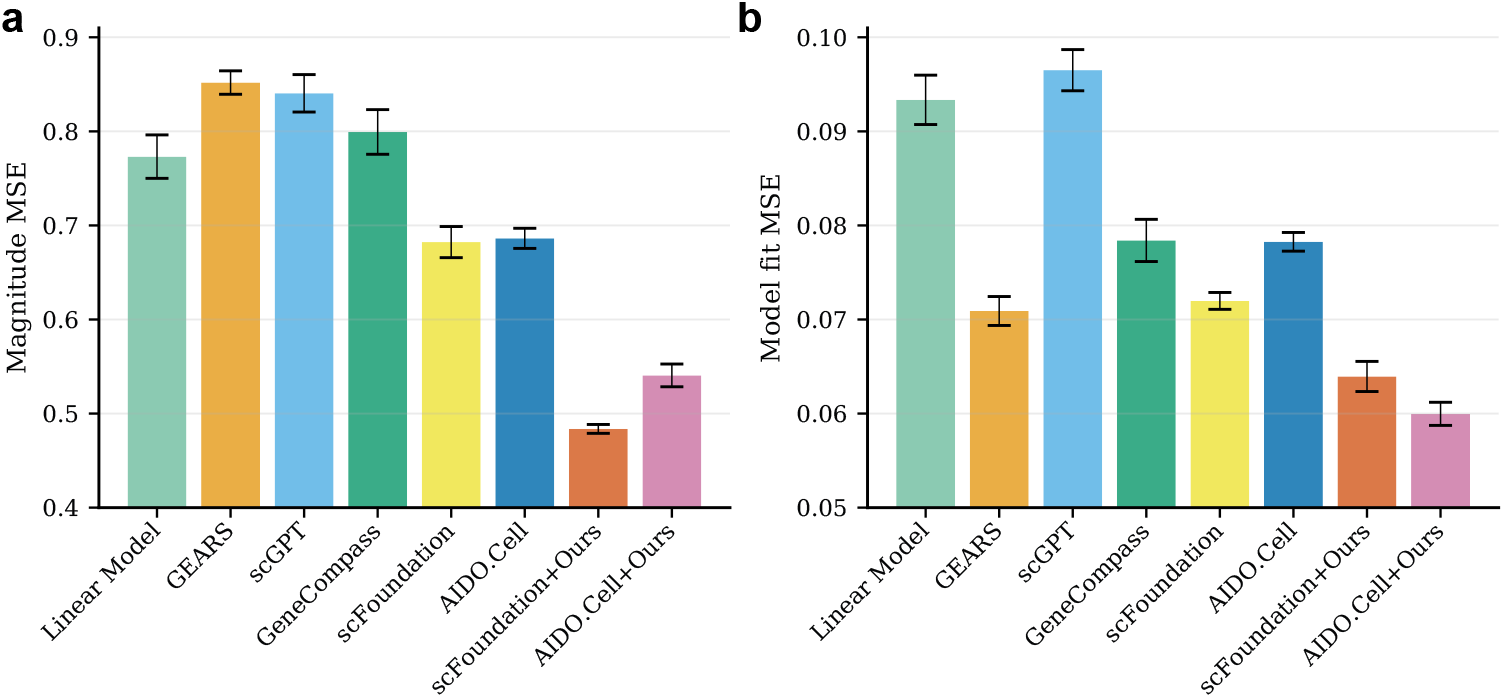
Evaluation of (a) magnitude and (b) model fit predictions for double-gene perturbations on the Norman dataset. We report the mean squared error (MSE) between the predicted and ground-truth magnitudes and model fits. The error bar represents the standard deviations of each model’s performance across 5 splits.

In summary, PertAdapt demonstrates superior accuracy in predicting double-gene perturbation effects, outperforming existing approaches across almost all evaluation settings. Its consistent gains in both in-distribution and OOD scenarios highlight its stronger generalization ability and its robustness in modeling combinatorial perturbational responses.

### 3.2 Generalizing single-gene perturbations across data regimes

PertAdapt delivers consistently strong performance across a range of single-gene perturbation datasets, as shown in Table 2. For single-gene perturbations, there is only one test scenario, seen 0/1, where the perturbed gene of the test set never appears in the perturbed genes of the training set. Across the six evaluated datasets, PertAdapt consistently achieves lower mean squared error (MSE) and higher Pearson correlation than most prior baselines. PertAdapt with either backbone foundation model (FM) improves the MSE by 17% on the Replogle RPE1 dataset, 28% on the Replogle K562 dataset, and by more than 30% on the remaining four datasets compared to the best-performing baselines. For the Pearson metric, PertAdapt with either backbone also achieves the highest scores on all datasets except for Replogle RPE1, where scFoundation alone reaches 0.731, slightly higher than AIDO.Cell+PertAdapt (0.725) but still below scFoundation+PertAdapt (0.737). Consistent with the double-gene results, sc-Foundation+PertAdapt performs slightly better and more stably than AIDO.Cell+PertAdapt in general. Among the six single-gene perturbation datasets, four (RPE1, K562, HepG2, and Jurkat) are sufficiently sampled, each containing more than 1,000 perturbations, whereas the remaining two (Adamson and Dixit) are smaller K562-based datasets comprising only 78 and 20 perturbations, respectively (Appendix Section A). The consistent improvement of PertAdapt across these diverse cell lines demonstrates its strong generalizability beyond specific experimental systems. Remarkably, PertAdapt also performs robustly under limited-data regimes, substantially outperforming all prior baselines on the smallest Dixit dataset (achieving *>* 90% reduction in MSE). This suggests that the perturbation adapter can effectively leverage the pretrained foundation model to capture transferable representations even when training samples are scarce.

**Table 2:** Evaluation of PertAdapt and baselines on the single-gene perturbation datasets. The datasets include Replogle RPE1, Replogle K562, Nadig HepG2, Nadig Jurkat, Adamson, and Dixit. We report results on the overall scenario (all tested perturbations), which is also the seen 0/1 scenario for single-gene perturbations

| Model | Replogle RPE1 |  | Replogle K562 |  | Nadig HepG2 |  |
| --- | --- | --- | --- | --- | --- | --- |
|  | MSE↓ | Pearson↑ | MSE↓ | Pearson↑ | MSE↓ | Pearson↑ |
| No-Change | 0.201 | 0.706 | 0.193 | 0.908 | 0.121 | 0.681 |
| Linear Model | 0.113 | 0.704 | 0.165 | 0.924 | 0.101 | 0.651 |
| GEARS | 0.159 | 0.719 | 0.124 | 0.929 | 0.101 | 0.652 |
| scGPT | 0.158 | 0.721 | 0.135 | 0.919 | 0.110 | 0.676 |
| GeneCompass | 0.170 | 0.727 | 0.121 | 0.921 | 0.104 | 0.680 |
| scFoundation | 0.166 | <u>0.731</u> | 0.142 | 0.927 | 0.109 | 0.684 |
| AIDO.Cell | 0.134 | 0.724 | 0.130 | 0.928 | 0.102 | 0.680 |
| scFoundation+Ours | **0.091** | **0.737** | **0.083** | <u>0.935</u> | **0.062** | **0.700** |
| AIDO.Cell+Ours | <u>0.094</u> | 0.725 | <u>0.088</u> | **0.936** | <u>0.067</u> | <u>0.685</u> |
|  | Nadig Jurkat |  | Adamson |  | Dixit |  |
|  | MSE↓ | Pearson↑ | MSE↓ | Pearson↑ | MSE↓ | Pearson↑ |
| No-Change | 0.111 | 0.801 | 0.905 | 0.787 | 0.380 | 0.831 |
| Linear Model | 0.101 | 0.815 | 0.675 | 0.862 | 0.229 | 0.912 |
| GEARS | 0.095 | 0.817 | 1.028 | 0.795 | 0.271 | 0.896 |
| scGPT | 0.112 | 0.806 | 1.116 | 0.679 | 0.295 | 0.841 |
| GeneCompass | 0.099 | 0.808 | 0.986 | 0.808 | 0.257 | 0.890 |
| scFoundation | 0.131 | 0.812 | 0.792 | 0.806 | 0.243 | 0.913 |
| AIDO.Cell | 0.103 | 0.820 | 0.804 | 0.840 | 0.213 | 0.924 |
| scFoundation+Ours | **0.061** | **0.826** | **0.324** | **0.904** | <u>0.021</u> | 0.992 |
| AIDO.Cell+Ours | <u>0.069</u> | <u>0.824</u> | <u>0.408</u> | <u>0.886</u> | **0.013** | **0.995** |

Interestingly, the simple linear model performs competitively with most nonlinear baselines, including the pretrained foundation models, across several datasets. This observation suggests that, without targeted adaptation, the expressive capacity of large foundation models may not be fully aligned with the perturbation-prediction objective, leading to performance that is effectively linear in nature. In contrast, PertAdapt substantially surpasses the linear baseline as well as the unadapted foundation models, indicating that the perturbation adapter effectively bridges the pretrained representations and the downstream perturbational response space. These results highlight the importance of task-specific adaptation in unlocking the potential of foundation models for single-cell perturbation modeling.

### 3.3 Scaling behavior with backbone FM size

To analyze the scaling trend of FM-based methods on genetic perturbation modeling, we evaluated the performance of PertAdapt using pretrained AIDO.Cell backbones of different sizes (3M, 10M, and 100M parameters) on the Norman and Replogle RPE1 datasets (Fig. 4). We further compared these results with the native AIDO.Cell method applied to each backbone size. Interestingly, while the native backbones exhibit a clear scaling trend (3M < 10M < 100M), PertAdapt maintains consistently strong performance across all three scales. This suggests that PertAdapt largely decouples perturbation modeling from backbone capacity: once the backbone provides a sufficiently rich basal representation (even at 3M), PertAdapt effectively captures perturbation-induced transcriptional shifts without requiring additional model capacity. In contrast, native methods depend directly on backbone expressiveness, resulting in a stronger scaling effect. While three publicly available model sizes do not allow us to claim a full scaling law, the observed monotonic trend remains consistent and informative, and we conclude that PertAdapt achieves robust and capacity-insensitive performance across backbones of different scales.

**Figure 4:**
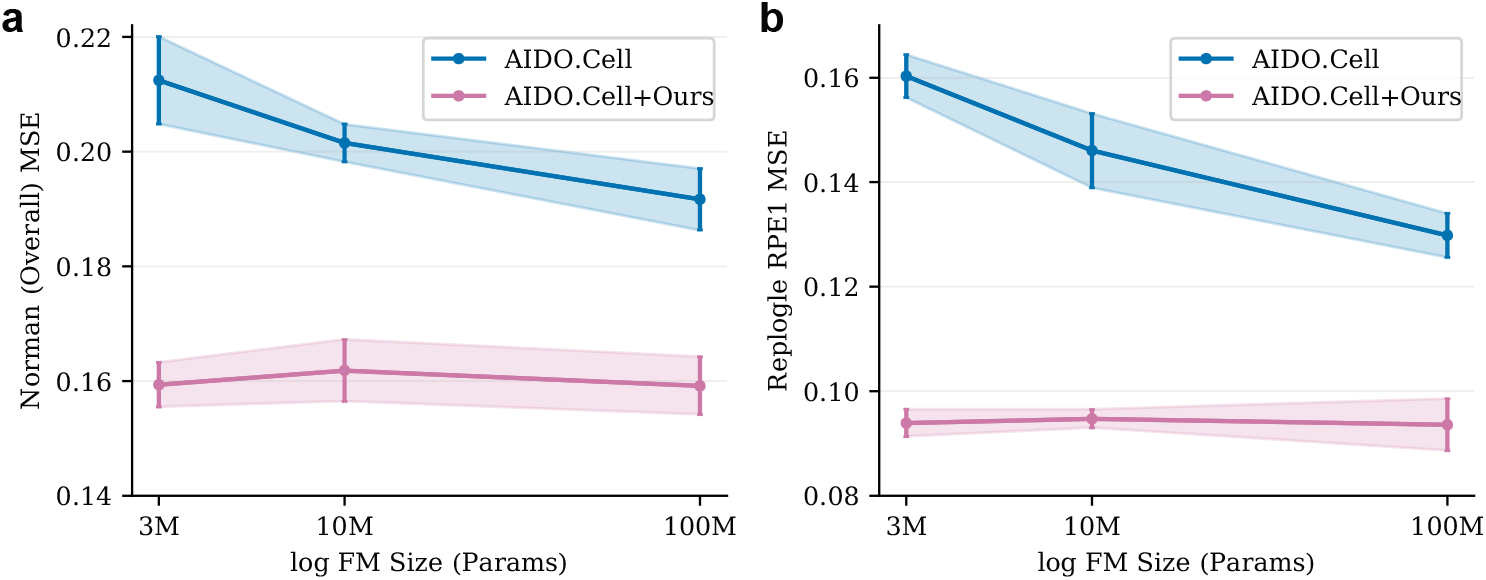
Scaling trend of perturbation-prediction MSE with respect to the size of the foundation model (FM). We compare AIDO.Cell and AIDO.Cell+Ours across FMs of different parameter magnitudes (3M, 10M, and 100M) on (**a**) the Norman dataset (overall performances) and (**b**) the Replogle RPE1 dataset. The x-axis is plotted on a logarithmic scale, and the error bar represents the standard deviations of each model’s performance across 5 splits.

### 3.4 Ablation study

The ablation results for both backbones of our method on the Norman (overall performances) and Replogle RPE1 datasets are presented in Table 3. We first assessed the contributions of the two main innovations of our method: the perturbation adapter and the adaptive loss. The configurations *FM+PA+AM* and *FM*+ ℒ_*adapt*_ reveal the performance gains of the adapter and the adaptive loss, respectively. In addition, as a key design within the adapter, the attention mask derived from gene functional similarities was evaluated independently to quantify its standalone effect. The improvement of *FM+PA+AM* over *FM+PA* confirms that the attention mask is an essential component of the perturbation adapter. We further verify the necessity of the pretrained FM by replacing it with a randomized counterpart. Overall, the results in Table 3 indicate that each component contributes positively to model performance, yet none alone achieves the effectiveness of the complete PertAdapt model that integrates all components. This highlights the synergistic advantage of jointly combining the pretrained FM, the adapter with the gene-similarity attention mask, and the adaptive loss in a unified framework.

**Table 3:**
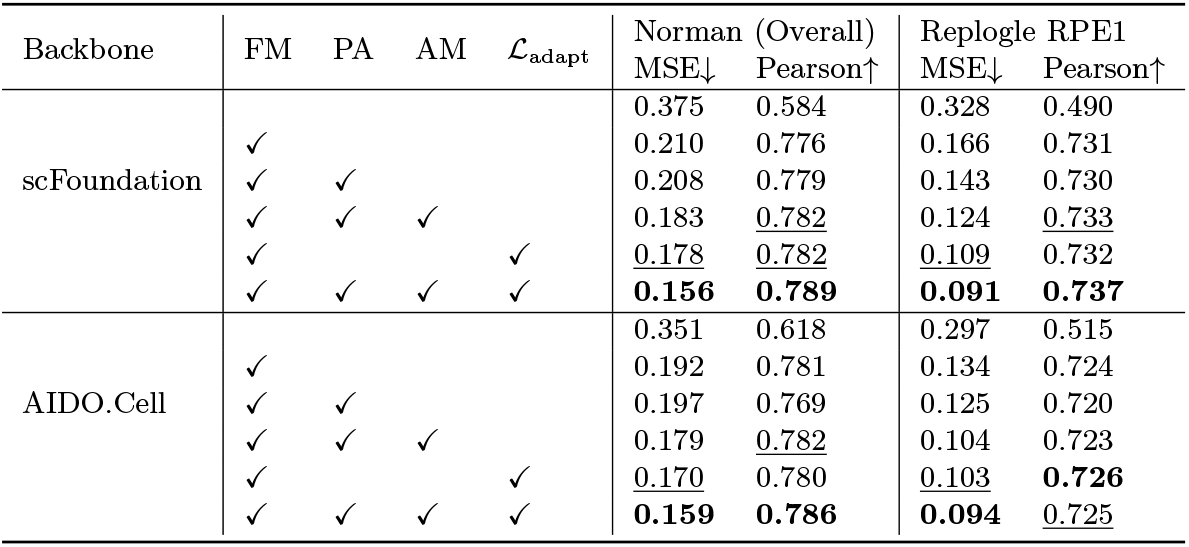
Ablation study of PertAdapt. We conduct ablation experiments using both backbones, and evaluate the contribution of four key components in PertAdapt: (1) the pretrained foundation model (FM); (2) the perturbation adapter (PA); (3) the adapter attention mask (AM); and (4) the adaptive loss (ℒ_adapt_). For each ablated variant, the removed component is replaced with: a randomized frozen FM, the backbone’s original decoder for perturbation, no mask, or the backbone’s original loss function for perturbation, respectively. Results are reported on the Norman dataset (overall performances) and the Replogle RPE1 dataset.

## 4 Conclusion and Limitations

In this work, we introduced PertAdapt, a plug-in framework that augments single-cell foundation models with a conditional adapter and an adaptive loss to better predict transcriptional responses to genetic perturbations. By integrating gene-similarity-masked attention and dynamically prioritizing perturbation-sensitive signals, PertAdapt enables more effective transfer of pretrained knowledge to perturbation tasks. PertAdapt consistently outperforms state-of-the-art methods in comprehensive experiments across seven diverse datasets of perturbations. PertAdapt shows the ability to capture multiplexed gene interaction behaviors in double-gene perturbation datasets, while exhibiting strong generalization in datasets with limited single-gene perturbations. Furthermore, a comparison of the scaling trend reveals its robustness in different backbone capacities. Ablation studies further validate the contribution of each component, demonstrating that both the perturbation adapter and the adaptive loss are essential for unlocking the predictive capacity of pretrained single-cell FMs.

Despite the clear performance gains achieved by PertAdapt, several limitations remain. First, its computational cost remains comparable to that of the native backbone methods. Future work may explore more parameter-efficient adapter designs to further reduce time and memory complexity. Second, although the GO mask strengthens biologically grounded relations, it may limit the model from capturing novel or previously unannotated interactions, which might be alleviated by combining GO with additional gene-similarity priors, e.g. PPI networks. Third, beyond genetic perturbations in single-cell data, PertAdapt may also be extended to broader perturbation modalities such as drugs and cytokines, as well as multi-omic integration and mixed perturbation spaces. We anticipate that PertAdapt will serve as a flexible foundation for building scalable and generalizable models in perturbation biology.

## A Dataset Details

We used three perturbation-effect datasets (Norman, Adamson, and Dixit) [2, 37, 1] preprocessed by scFoundation [19], together with four additional datasets (Replogle RPE1, Replogle K562, Nadig HepG2, Nadig Jurkat) [6, 7] that we preprocessed using the same procedure. In preprocessing, we first normalized the raw count matrices using log1p, then retained the 19,264 genes drawn from the input gene list used by scFoundation and AIDO.Cell [23], and finally filtered low-quality cell samples and conditions, keeping cell samples with at least 200 nonzero genes and perturbation conditions with at least 30 samples. Detailed statistics for all datasets after preprocessing are listed in Appendix Table A1. In addition, the distribution of per-gene counts of genes sharing *≥* 1 Gene Ontology (GO) terms is in Fig. A1.

**Table A1:** Details of the preprocess perturbational effects datasets. Here # means the number of something. The Norman dataset is a two-gene perturbation dataset [2]. The other 6 datasets are single-gene perturbation datasets [37, 1, 6, 7].

| Dataset | # 1-gene perturbations | # 2-gene perturbations | # Cells | # 1-gene perturbed cells | # 2-gene perturbed cells |
| --- | --- | --- | --- | --- | --- |
| Norman | 101 | 125 | 84,986 | 45,342 | 32,369 |
| Replogle RPE1 | 1,543 | 0 | 162,733 | 151,248 | 0 |
| Replogle K562 | 1,092 | 0 | 162,751 | 152,060 | 0 |
| Nadig HepG2 | 1,615 | 0 | 118,877 | 113,901 | 0 |
| Nadig K562 | 1,580 | 0 | 141,428 | 132,904 | 0 |
| Adamson | 78 | 0 | 63,857 | 44,113 | 0 |
| Dixit | 20 | 0 | 37,160 | 34,086 | 0 |

**Figure A1:**
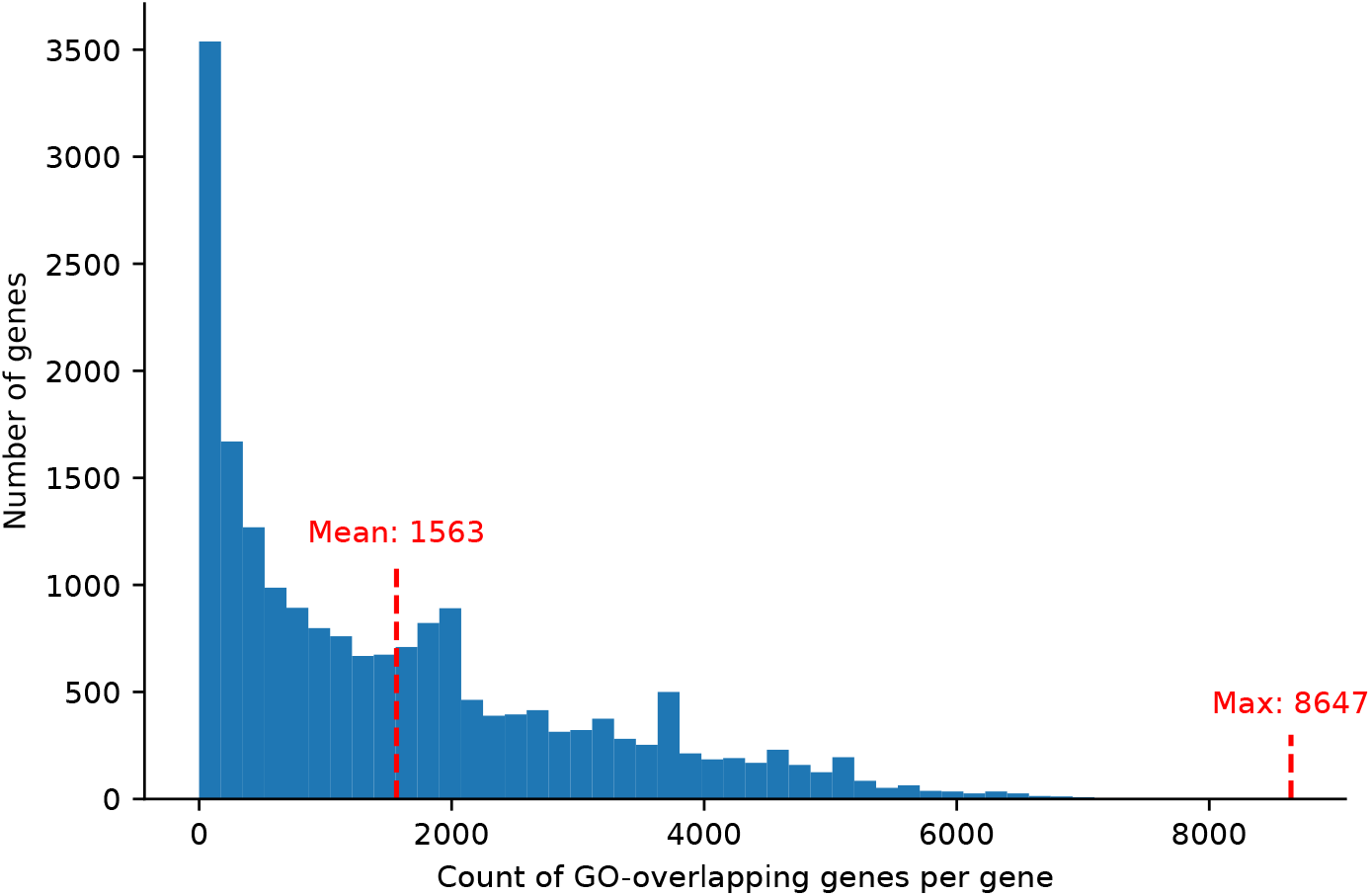
Histogram of per-gene counts of genes sharing at least one Gene Ontology (GO) term. Each bar represents the number of genes that share a given number of GO-overlapping genes, illustrating the distribution of sparsity in the GO-based attention mask. The mean and maximum numbers of GO-overlapping genes per gene are 1, 563 and 8, 647, respectively, as indicated above the red dashed lines.

## B Evaluation Metrics Details

Mean Squared Error (MSE) and Pearson correlation coefficient (Pearson) were employed to evaluate the model’s performance. For pseudobulk expressions, i.e. the mean expression of each gene across all cells under the same perturbation condition, **ŷ** and ground-truth **y** under perturbation condition *c* of *k* = 1, 2, …*N* genes, the MSE and Pearson on a given set of conditions 𝒞 and a set of genes 𝒟_*c*_ are:

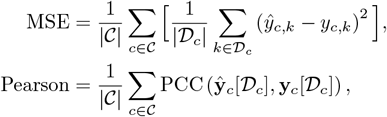

where PCC(·, ·) denotes the Pearson correlation coefficient of 2 vectors, and the vector **y**[𝒟] denotes the subvector of **y** indexed by 𝒟.

In our evaluations, the set of conditions 𝒞 varies according to the experimental setup. When defined on the Norman dataset, 𝒞 includes all tested perturbations for the *Overall* results, and is further divided into *Seen 0/2, Seen 1/2*, and *Seen 2/2* subsets. For all single-gene perturbation datasets, the only 𝒞 is the *Seen 0/1* results, which includes all the tested perturbations. The set of genes 𝒟_*c*_ varies from subsets containing different numbers of differentially expressed (DE) genes given the perturbation condition *c* to the set of all genes.

In classifying gene interaction types of double-gene perturbations, we employ the magnitude and model fit scores following [2, 14]. Let **x** denote the average expression vector of all unperturbed cells. The difference **y** − **x** represents the perturbation effect of a double-gene perturbation *c* = {*i, j*}, denoted as **Δ**_*i*,*j*_. Similarly, we compute the average perturbation effects **Δ**_*i*_ and **Δ**_*j*_ for single-gene perturbations *c* = {*i*} and *c* = {*j*}, respectively. We then solve the following linear model:

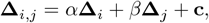

where *α* and *β* are scalars representing the linear combination coefficients, and **c** is an offset vector. The model is fit using robust regression with a Theil-Sen estimator. Let the estimated coefficients be 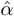 and 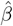. The magnitude and model fit scores are then defined as

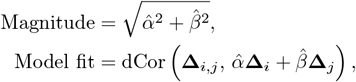

where dCor(·, ·) denotes the distance correlation coefficient between two vectors. For each model, the magnitude and model fit scores are evaluated by replacing **y** with its predicted counterpart ŷ. We then compute the mean squared error (MSE) between these predicted scores and their ground-truth counterparts, as reported in Main Fig. 3.

## C Hyperparameter Tuning

We selected the optimal set of hyperparameters via the validation process on the Norman dataset [2]. The search was conducted over the following ranges: the DE size of the adaptive loss *k* ∈ {20, 50, 100, 200}, the embedding dimension *D* ∈ {64, 256, 512, 1024}, the number of attention heads *H* ∈ {2, 4, 8, 16}, and the head dimension *d*_*h*_ = *D/H* within the perturbation adapter. The effective batch size was fixed at 32 to fully utilize the available GPUs, and the Adam optimizer was employed with a default learning rate of 10^−3^. The best-performing configuration was found to be *D* = 512 and *H* = 8.

The backbone perturbation encoder is the GNN module adopted from the GEARS model [14]. Its hidden dimension is fixed to 512, matching that of our model, while the number of GNN layers remains the same as in the original design (1 layer). The pretrained backbone foundation models (FMs) are structured as follows. For scFoundation, which contains approximately 100 million parameters, the architecture consists of a Transformer encoder with 12 layers, 12 attention heads, and a hidden size of 768, followed by a Performer decoder with 6 layers, 8 heads, and a hidden size of 512. For AIDO.Cell, a Transformer decoder-only model, three parameter scales are used: 3M (6 layers, 4 heads, 128 hidden size), 10M (8 layers, 8 heads, 256 hidden size), and 100M (18 layers, 20 heads, 640 hidden size). Both backbone models encode the full expression profiles of 19,264 genes.

## D Experimental Results on different sets of genes

We evaluate the model performance using mean squared error (MSE) and Pearson correlation scores on the top 20, 50, 100, and 200 differentially expressed (DE) genes, as well as on all genes. While the top 20 DE results are presented in the main manuscript, the results for other gene sets are provided in Table A2. Across all evaluation settings, PertAdapt consistently outperforms most baseline methods on both metrics, demonstrating its robustness across different subsets of DE genes and the full gene set. Notably, expanding from the top-20 DE genes to larger sets introduces many perturbation-insensitive genes, causing a marginal improvement. Hence, the comparison between the top 20 DE genes concentrates informative signal and better separates models.

**Table A2:** Evaluation of PertAdapt and baselines on the Norman dataset. The performances over all tested perturbations on different numbers of DE genes and all genes are presented.

| Model | 50 DE genes |  | 100 DE genes |  | 200 DE genes |  | All genes |  |
| --- | --- | --- | --- | --- | --- | --- | --- | --- |
|  | MSE↓ | Pearson↑ | MSE×0.1 ↓ | Pearson↑ | MSE×0.1 ↓ | Pearson↑ | MSE×0.01 ↓ | Pearson↑ |
| No-Change | 0.213 | 0.796 | 1.407 | 0.865 | 0.950 | 0.908 | 0.340 | 0.988 |
| Linear Model | 0.133 | <u>0.837</u> | 0.820 | 0.889 | 0.534 | <u>0.925</u> | 0.238 | <u>0.992</u> |
| GEARS | 0.129 | 0.830 | 0.811 | 0.879 | 0.533 | 0.920 | 0.291 | 0.990 |
| scGPT | 0.119 | 0.824 | 0.801 | 0.873 | 0.575 | 0.908 | 0.299 | 0.984 |
| GeneCompass | 0.117 | 0.831 | 0.719 | 0.869 | 0.573 | 0.907 | 0.286 | 0.959 |
| scFoundation | 0.107 | 0.820 | 0.687 | 0.884 | 0.561 | 0.921 | 0.264 | 0.983 |
| AIDO.Cell | 0.112 | 0.836 | 0.626 | 0.877 | 0.532 | 0.909 | <u>0.236</u> | 0.986 |
| scFoundation+Ours | **0.094** | **0.846** | <u>0.604</u> | **0.895** | **0.498** | **0.928** | **0.235** | **0.993** |
| AIDO.Cell+Ours | <u>0.103</u> | **0.846** | **0.600** | <u>0.892</u> | <u>0.510</u> | <u>0.925</u> | <u>0.236</u> | 0.989 |

